# Improved Score Statistics for Meta-analysis in Single-variant and Gene-level Association Studies

**DOI:** 10.1101/195545

**Authors:** Jingjing Yang, Sai Chen, Gonçalo Abecasis, IAMDGC

**Author notes:** Correspondence authors: J.Y. and G.A.

## Abstract

Meta-analysis is now an essential tool for genetic association studies, allowing these to combine large studies and greatly accelerating the pace of genetic discovery. Although the standard meta-analysis methods perform equivalently as the more cumbersome joint analysis under ideal settings, they result in substantial power loss under unbalanced settings with various case-control ratios. Here, we investigate why the standard meta-analysis methods lose power under unbalanced settings, and further propose a novel meta-analysis method that performs as efficiently as joint analysis under general settings. Our proposed method can accurately approximate the score statistics obtainable by joint analysis, for both linear and logistic regression models, with and without covariates. In addition, we propose a novel approach to adjust for population stratification by correcting for known population structures through minor allele frequencies (MAFs). In the simulated gene-level association studies under unbalanced settings, our method recovered up to 85% power loss caused by the standard method. We further showed the power gain of our method in gene-level association studies with 26 unbalanced real studies of Age-related Macular Degeneration (AMD). In addition, we took the meta-analysis of three studies of type 2 diabetes (T2D) as an example to discuss the challenges of meta-analyzing multi-ethnic samples. In summary, we propose improved single-variant score statistics in meta-analysis, requiring “accurate” population-specific MAFs for multi-ethnic studies. These improved score statistics can be used to construct both single-variant and gene-level association studies, providing a useful framework for ensuring well-powered, convenient, cross-study analyses.

## Introduction

Meta-analysis is now an essential tool for genetic association studies, allowing these to combine information on 100,000s – 1,000,000s of samples, and greatly accelerating the pace of genetic discovery. Under ideal experiment settings, e.g., the same case-control ratio for all individual studies, the standard meta-analysis methods perform as efficiently as the more cumbersome alternative of sharing individual-level data^1^. Standard meta-analysis methods have been routinely used in many large-scale genome-wide association studies (GWASs), identifying many complex trait loci, e.g., type 2 diabetes (T2D)^2^-^4^, lipid levels^5^, body mass index (BMI)^6^, rheumatoid arthritis^7^, and fasting glucose levels^8^. Many tools implementing standard meta-analysis methods have been proposed for both single-variant and gene-level association studies, such as METAL for single-variant association studies^9^, META-SKAT, MASS, and RAREMETAL for gene-level association studies^10^-^13^ However, when the case-control ratios (or phenotype means and variances for quantitative traits) vary among individual studies due to unbalanced study designs (e.g., studies using Biobank^14^ data), the current standard meta-analysis methods result in substantial power loss. The limitations of current standard methods apply where meta-analysis is based on combining sample sizes and p values (the Stouffer method^15^), using regression coefficients and their standard errors (the Cochran method^16^), or by combining score statistics^10^. Although weighting by effective sample sizes of individual studies may improve the power for single-variant meta-analysis using the Stouffer and Cochran methods^9^, the weighting strategy will fail for gene-level meta-analysis based on score statistics. This is because the magnitudes of score statistics are in the order of sample sizes (unlike the unit-free test statistics used by the Stouffer and Cochran methods). We note that the standard meta-analysis methods based on score statistics^10; 13^ ignores the between-study variances of both phenotype and genotype by directly summing within-study score statistics to approximate the “joint” score statistics. This will cause power loss when the between-study variances contain association information as in the scenario of unbalanced studies.

Here, we describe a novel meta-analysis method that models the between-study variances, thus improving power under unbalanced settings. Our method is suitable for both linear and logistic regression models, with and without covariates. When the study samples are of the same population (i.e., without population stratification), our method is equivalent to the more cumbersome joint analysis (i.e., golden standards). For studies with population stratification, where joint analysis is expected to cause inflated false positive rates, we propose a novel approach to adjust for population stratification using known population structure information, i.e., population-specific minor allele frequencies (MAFs). Observing that population stratification is reflected by MAFs in the score statistics, we adjust population stratification by regressing out the effects of population-specific MAFs (obtainable from reference panels such as 1000 Genome^17^, or Biobanks^14^) from within-study MAFs. This approach aims to adjust for the between-study variations due to population stratification, which can be implemented as a complementary step of adjusting for first few principle components (PCs) within individual studies^18^ (adjusting for the within-study population stratification). Although our approach applies to any meta-analysis methods based on score statistics, we focus on single-variant score tests^19^, and further extend to enable gene-level Burden^20,21^; test and sequential kernel association test (SKAT)^22^.

By simulation studies, we showed that, under unbalanced settings, our method recovered up to 84% power loss caused by the standard methods while controlling for false positive rates (i.e., type I errors), regardless of the existence of population stratification. Further, we demonstrated the power gain of our method in the real gene-level association studies of Age-related Macular Degeneration (AMD)^23^, consisted of 26 unbalanced individual studies and 33,976 unrelated European samples (Table S1). For example, the known AMD risk gene *CFI* has SKAT p value 1.9 × 10^−10^ by joint analysis, p value 1.2 × 10^−4^ by the standard meta-SKAT, and p value 3.1 × 10^−9^ by our meta-SKAT. In addition, we applied our method on the meta-analysis of three studies of type 2 diabetes (T2D) with Finnish and American European populations. We noticed that the standard methods were desirable when the between-study variances contain non-association variances that can not be completely corrected for, such as when individual studies use different metrics for phenotypes and covariates, or when the population-specific MAFs are unavailable or “inaccurate”.

In summary, we propose improved single-variant score statistics for meta-analysis to achieve the same performance as the joint analysis under general settings. These improved single-variant score statistics can be used for both single-variant association studies and combined to conduct gene-level association studies. Our method provides a useful framework for ensuring well-powered, convenient, cross-study analyses and is now implemented in the RAREMETAL software.

## Material and Methods

### Score Statistics for Individual Studies

Consider conducting meta-analysis with *K* studies, where the. *k*th study has *n*_*k*_ samples genotyped at *m*_*k*_ variants. Let *y*_*k*_ denote the *n*_*k*_ × 1 phenotype vector; X_*k*_ denote the *n*_*k*_ × *m*_*k*_ genotype matrix, encoding the minor allele count per individual per variant as (0, 1, 2); and *C*_*k*_ denote the n_*k*_ × (*q*_*k*_ + 1) augmented covariate matrix with the first column set to 1’s and the others encoding *q_k_* covariates. For each individual study, we consider the standard linear regression model (Equation 1) for quantitative traits

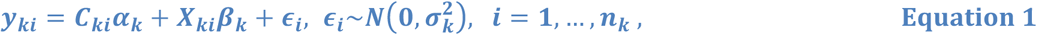

and the standard logistic regression model (Equation 2) for dichotomous traits

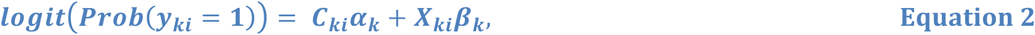

where *X*_*ki*_ is the *i*th row of genotype matrix *X*_*k*_, *β*_*k*_ is the vector of genetic effect-sizes, *C*_*ki*_ is the *i*th row of augmented covariate matrix *C*_*k*_, and *α*_*k*_ is the vector of covariate effects including the intercept term. Let **u**_*k*_ denote the vector of score statistics for the *k*th study and **V**_*k*_ denote the variance-covariance matrix of **u**_*k*_ (Appendix A).

### Standard Meta-analysis

For notation simplicity, we assume all-studies measure the same set of variants, with phenotypes from the same underlying distribution. The current standard meta-analysis methods typically approximate the joint score statistics (obtainable by joint analysis) by

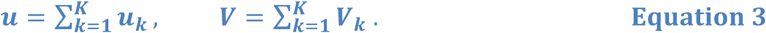

Under unbalanced studies, these statistics will be systematically different from those obtained from the joint analysis using combined individual-level data – potentially leading to substantial power loss. Instead, we derive our improved approximations for these joint score statistics (*u, V*) directly from the statistic formulas using combined data as in joint analysis.

### Simplified Case without Covariates

First, we consider a simplified case without covariates, in which the following analytical formulas (Equations 4 and 5) are derived for joint score statistics under both linear and logistic regression models (Appendix B.1), in terms of the within-study score statistics (***u***_*k*_, ***V***_*k*_), sample size *n*_*k*_, phenotype mean deviation *δ*_*k*_, residual variance estimate 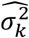, and MAF vector ***f***_*k*_,

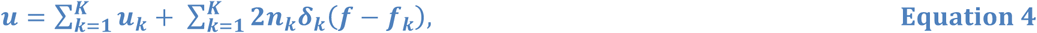

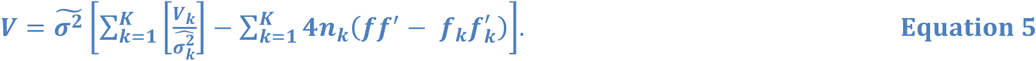

Here, 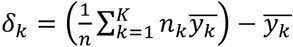 denotes the difference between the overall phenotype mean and within-study phenotype mean; 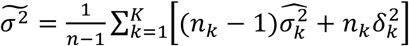 denotes the joint residual variance; and 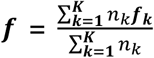 denotes the overall MAF vector. The key difference from the standard approach (Equation 3) is that we actually model the between-study variations (second terms in Equations 4 and 5) through the phenotype mean deviation **δ**_*k*_, residual variance difference between 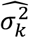and 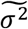, and MAF difference between ***f***_*k*_ and ***f***.

We note that, when the studies are balanced with samples of the same population, i.e., 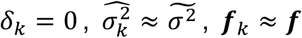, Equations 4 and 5 are equivalent to the standard estimates in Equation 3 and equivalent to the joint estimates. This is why the standard meta-analysis methods perform as efficient as joint analysis in balanced studies with samples of the same population. However, when the studies are unbalanced (e.g., with case-control ratios differ greatly among studies), i.e., 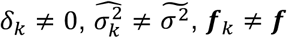, the standard estimates (Equation 3) can no longer accurately approximate the joint score statistics, potentially leading to substantial power loss. In contrast, our meta-analysis method (using Equations 4 and 5) will be equivalent to joint analysis under general settings.

### General Case with Covariates

Second, we consider the general case with covariates in which the joint score statistic ***u*** is still given by Equation 4, but the formula for the joint variance-covariance matrix ***v*** will be different from Equation 5. In this case, we approximate the phenotype mean deviation 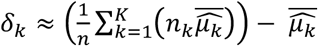, where 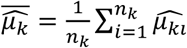 is the average of the fitted phenotypes in study *k* under the null regression models with *β* = 0 (Equations 1 and 2).

For notation simplicity, we assume all individual studies have the same set of covariates. Then under the linear regression model (Equation 1), we can estimate ***V*** by

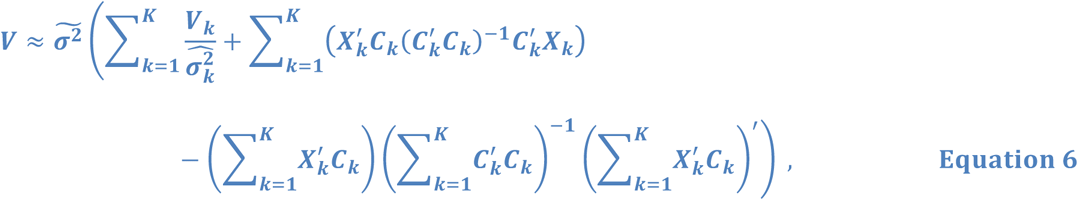

where the quantities of the covariate relationship matrix 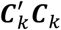 and genotype-covariate relationship matrix 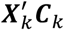 need to be shared (Appendix B.2).

Under the logistic regression model (Equation 2), T can be estimated by

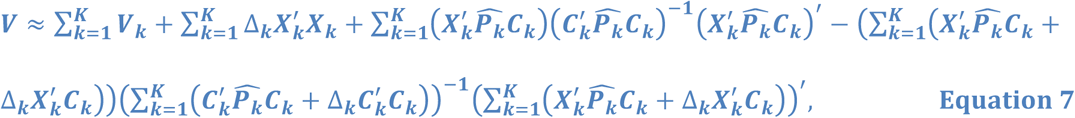

where 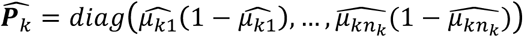 denotes the diagonal matrix of phenotypic variances after correcting for within-study covariates; 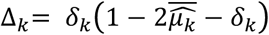 is the average difference between 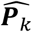 and an analogous estimate in joint analysis. To enable the calculation by Equation 7, the quantities of the genotype relation matrix 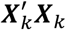, covariate relation matrices 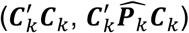, and the genotype-covariate relation matrices 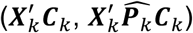 need to be shared (Appendix B.2).

### Adjusting for Population Stratification

Because our score statistic formulas (Equations 4-7) are equivalent to the analogous statistics obtainable in joint analysis, the meta-analysis using our score statistic estimates is likely to subject to inflated false positive rates (as joint analysis) when studies are of multi-ethnic. We note that the population stratification is reflected by the between – study variances, particularly by the differences between within-study MAFs and the joint MAFs, i.e., (***f**-**f**_***k***_*) in Equation 4 and 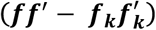 in Equation 5. The standard meta-analysis methods using Equation 3 fail to model the between-study variances (free of population stratification), for dropping the second terms in Equations 4 and 5. Therefore, when population stratification exists in the combined data, we have to adjust our score statistic estimates (by Equations 4 and 5) to correct for inflated false positive rates.

Here, we propose to normalize our within-study MAFs by regressing out the population effects that can be explained by known population-specific MAFs (e.g., population MAFs from reference panels like 1000 Genome Project^17^, or Biobanks^14^). For example, with known MAF vectors ***f**_EUR_, **f**_AMR_, f_AFR_, **f**_SAS_, **f**_EAS_* of genome-wide variants for European, American, African, South Asian, and East Asian populations in the 1000 Genome Project^17^, we first fit the following linear regression model per individual study using the MAFs of genome-wide variants in the reference panel

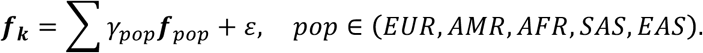

Then, in Equations 4 and 5, we substitute ***f***_***k***_ by the residuals 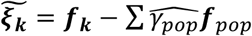, and ***f*** by the weighted residual averages 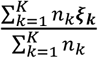. We set the corresponding elements in vectors ***f***_***k***_ and ***f*** as 0 for variants absent from the reference panel or with fitted values outside of the 95% predictive intervals, such that the between-study variances related to these variants will not be modeled in our method. Equivalently, in Equations 6 and 7, we can normalize the genotype matrix by 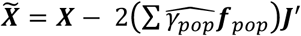 for variants in the reference panel, while set the genotype matrix as 0 for variants with unknown population-specific MAFs or with outlying fitted values.

### Practical Approach

Although Equations 6 and 7 enable joint-equivalent corrections for covariates in meta-analysis, they are not directly applicable in practice for the difficulties of sharing the quantities of 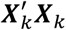, 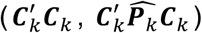, and 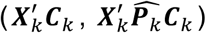 Thus, for computational simplicity, we suggest using Equation 5 with phenotypes corrected for covariates within individual studies under the linear regression model (Equation 1), where the dichotomous traits could be treated as quantitative traits by coding cases as 1’s and controls as 0’s. The RAREMETAL software also implements this practical approach. Both approaches (Equations 6 and 7 vs. Equation 5) produced nearly the same association results in our simulations. For both quantitative and dichotomous studies in this paper, we first corrected phenotypes within studies, and then used score statistic estimates by Equations 4 and 5 for association studies (adjusting for possible population stratification).

One key point of conducting joint-equivalent meta-analysis by our method is to include the intercepts from covariate correction into the corrected phenotypes, for modeling the between-study variances due to phenotype mean deviations. Otherwise, the phenotype deviation δ_*k*_’s will all be 0’s, and our score statistic estimates (Equation 4) will be equivalent to the standard estimates (Equation 3). Another key point is to make sure phenotype deviation δ_*k*_ ’s contain no other artificial effects (e.g., batch effects, effects due to different metrics or different underlying distributions across studies for phenotypes), which is likely to inflate the false positive rates.

### Test Statistics

Our meta-analysis methods are based on accurately approximating the joint score statistics (***u***, ***V***), and properly adjusting for possible population stratification. We consider the Burden test^20^ with statistic 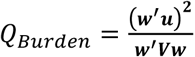 and SKAT^10^ with statistic 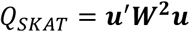 as examples of gene-level tests, where 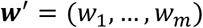 is the variant-specific weight vector, and ***W*** = *diag* (*w*_1_,…, *w*_*m*_ is the *m×m* diagonal matrix. For each variant, we take the weight as “capped” beta density value 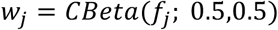 with the corresponding MAF ***f***_j_, to avoid assigning extremely large weights for extremely rare variants 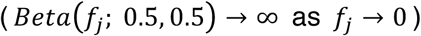. That is, with sample size *n*, 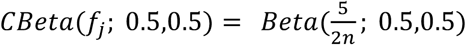 if the minor allele count 2*nf*_*j*_ < 5, otherwise 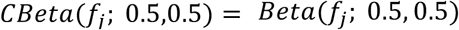 allowing equal variance contributions from all variants.

Under the null hypothesis 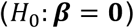, both Q_Burden_ and the the single-variant score test with statistic 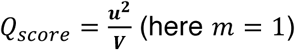 (here m = 1) follow a chi-square distribution with 1 degree of freedom (*df* = 1). Under the equivalent null hypothesis 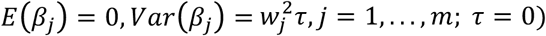 for SKAT, QSKAT asymptotically follows a mixture of chi-square distributions, 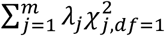, where 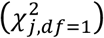 are independent chi-square random variables with *df* = 1, and *λ_j_*’s are nonzero eigenvalues of the variant relationship matrix ***∅ =WVW***.

### Simulation Studies

To evaluate the false positive rate (type I error) and power of our meta-analysis method, we conducted simulation studies in various scenarios with balanced and unbalanced studies, with and without population stratification, with quantitative and dichotomous traits (see details of the simulation setup in Appendix C).

Briefly, we first simulated haplotypes of three populations (European (EUR), Asian (ASA), and African (AFR)) by COSI with the well calibrated coalescent model^24^. Then we sampled genotypes of 1 × 10^5^ individuals per population with 339 variants, 96% of which have MAFs < 5%. Random risk regions of 100 variants were selected to simulate both quantitative and dichotomous phenotypes, according to the standard linear and logistic models. We simulated phenotypes under the null models (***β* = 0**) for evaluating the empirical type I error, and phenotypes with half of the variants in the risk regions as true causals for evaluating the power.

We considered meta-analysis with 5 individual studies and a total sample size 3,000 (Table S2), under combined scenarios of dichotomous and quantitative traits, balanced and unbalanced settings, common and uncommon covariates, single and multiple ethnic samples. For the balanced scenarios, each dichotomous study has 300 cases and 300 controls, while each quantitative study has 600 samples. For unbalanced dichotomous studies, there are (60, 180, 300, 420, 540) cases and (540, 420, 300, 180, 60) controls, such that the individual studies have the same sample size but various cases-control ratios. Unbalanced quantitative studies have sample sizes (200, 400, 600, 800, 1000). Two covariate scenarios were simulated: (i) common covariates for all studies; (ii) different covariates among studies.

For the case with single-ethnic samples (i.e., without population stratification), we compared our adjusted meta-analysis methods with the standard methods and the joint analysis, where the results by joint analysis will serve as the golden standards. For the case with multi-ethnic samples (i.e., with population stratification; with EUR samples in studies 1 and 3, ASA samples in studies 2 and 4, and AFR samples in study 5), we only considered balanced and unbalanced dichotomous studies with common covariates (Table S2). In this case, we corrected the population stratification using the population MAF vectors (*f*_EUR_, *f*_ASA_, *f*_AFR_) that were calculated from 1 × 10^5^ samples of the respective population. We compared our methods with the standard methods and joint analysis with first 4 principle components (PCs) as additional covariates.

### AMD and T2D Data

The study of age-related macular degeneration (AMD) by the International AMD Genomics Consortium (IAMDGC)^23^ consists of 26 individual studies with 33,976 European, 1,572 Asian, and 413 African unrelated samples. Variants were genotyped on a customized Exome-Chip and imputed against the 1000 Genome Project Phase I reference panel. Advanced AMD cases include both cases with choroidal neovascularization and cases with geographic atrophy^23;25^.

Three GWASs of type 2 diabetes (T2D) were considered in this paper: the Finland-United States Investigation of NIDDM genetics (FUSION) study^2^, METabolic Syndrome In Men (METSIM) study^26^, and Michigan Genomics Initiative (MGI) study. We analyzed 2,297 unrelated Finnish samples (1,142 cases vs. 1,155 controls) in FUSION, 3,340 unrelated Finnish samples (673 cases vs. 2,667 controls) in METSIM, and 16,495 unrelated European American samples (1,942 cases vs. 14,553 controls) in MGI.

For the association studies of both AMD and T2D, all participants gave informed consent and the University of Michigan IRB approved our analyses.

## Results

### Empirical Type I Errors in Simulation Studies

We repeated 2.5×10^7^ null simulations per scenario to obtain empirical type I errors with significance levels *α* = ^10-2^, ^10-4^, 2.5 × ^10-6^. We showed that both Burden test and SKAT –– by our adjusted meta-analysis method, the standard method, and joint analysis –– controlled well for type I errors in all scenarios without population stratification (Figure 1A; Figures S2 and S3).

**Figure 1.**
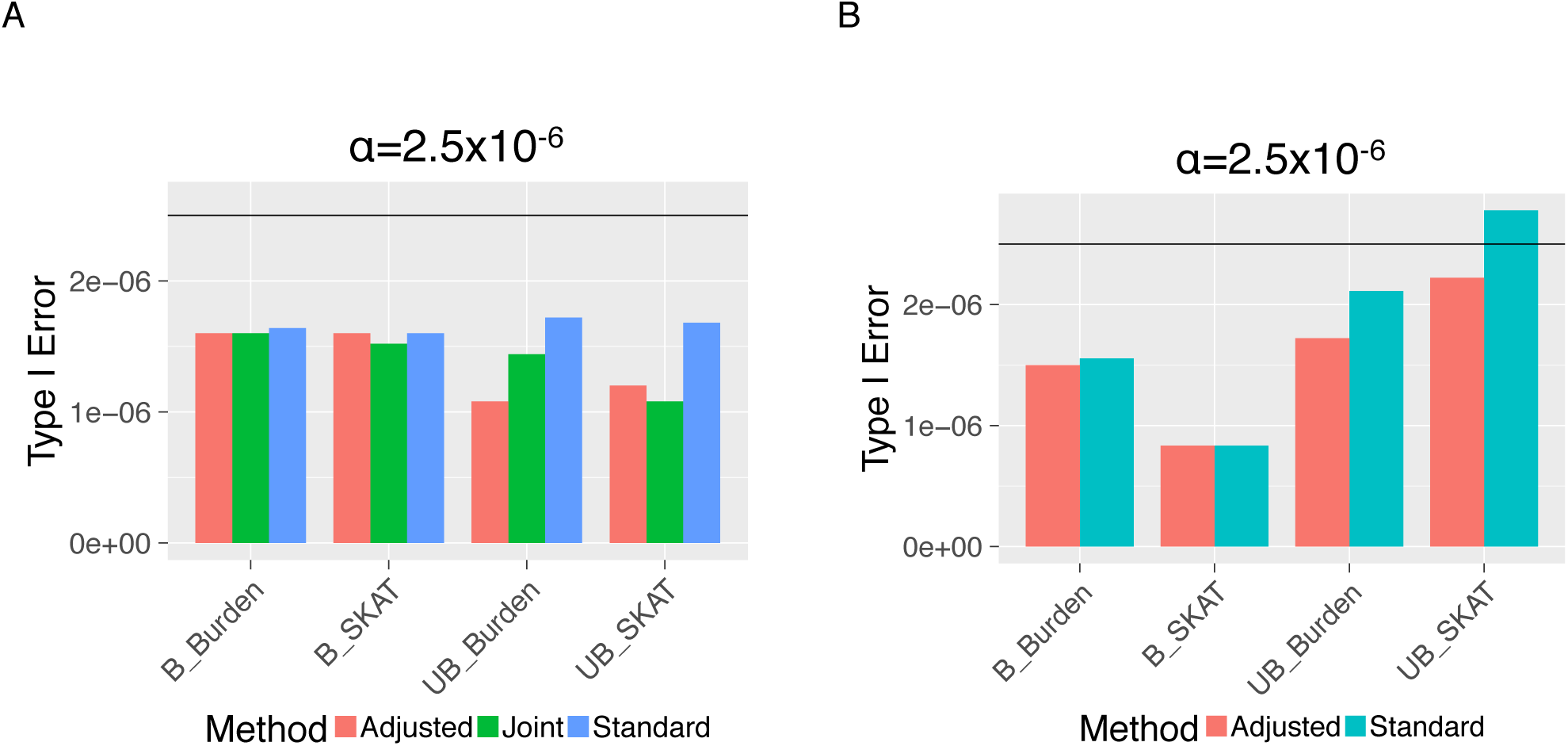
Empirical type I errors (with significance level α = 2.5 × 10^−6^) for null simulations of balanced (B) and unbalanced (UB) dichotomous studies, with common covariates. A: Scenario without population stratification; B: Scenario with population stratification. “Adjusted” denotes our new meta-analysis methods; “Standard” denotes the standard meta-analysis methods; and “Joint” denotes the joint analyses using combined individual-level data.

In the scenarios with population stratification, we showed that both Burden test and SKAT by our method and the standard method still controlled well for type I errors (Figure 1B; Figures S4 and S5 (C, D, E, F)), while the tests by joint analysis with first 4 joint PCs as additional covariates had highly inflated type I errors (see Quantile-Quantile (QQ) plots of – log10 (p values) in Figure S5 (A, B)). This demonstrated that the standard methods were not affected by population stratification, and that our approach of adjusting for population stratification successfully corrected for inflated type I errors.

### Empirical Power in Simulation Studies

For each scenario, we repeated 10,000 simulations to obtain the empirical power that is given by the proportion of simulations with p values < 2.5×10^−6^ (genome-wide significance level for gene-level association tests). Here, our goal is to compare the power of our adjusted meta-analysis method with the standard method and the joint analysis. The power differences between Burden test and SKAT will depend on simulation settings.

In the balanced dichotomous studies without population stratification, both standard and our adjusted estimates of joint score statistics (by Equation 4) were highly concordant with the golden standards obtained by joint analysis (R^2^ > 99.8%; Figure 2 (A, B)). In the unbalanced dichotomous studies, the standard meta estimates of score statistics (by Equation 3) scattered further from the joint score statistics (R^2^ ∽ 78.2%, Figure 2C), while our adjusted estimates were still concordant with the joint score statistics (R^2^ > 99.8%; Figure 2D). Consequently, under balanced settings, the p values of single-variant score tests by both standard methods and our adjusted methods were concordant with the joint analysis results (Figure 2(E, F)). However, under unbalanced settings, the p values by standard methods were less significant than joint analysis results (Figure 2G), hence less significant than the results by our adjusted methods that were concordant with the joint analysis results (Figure 2H).

**Figure 2.**
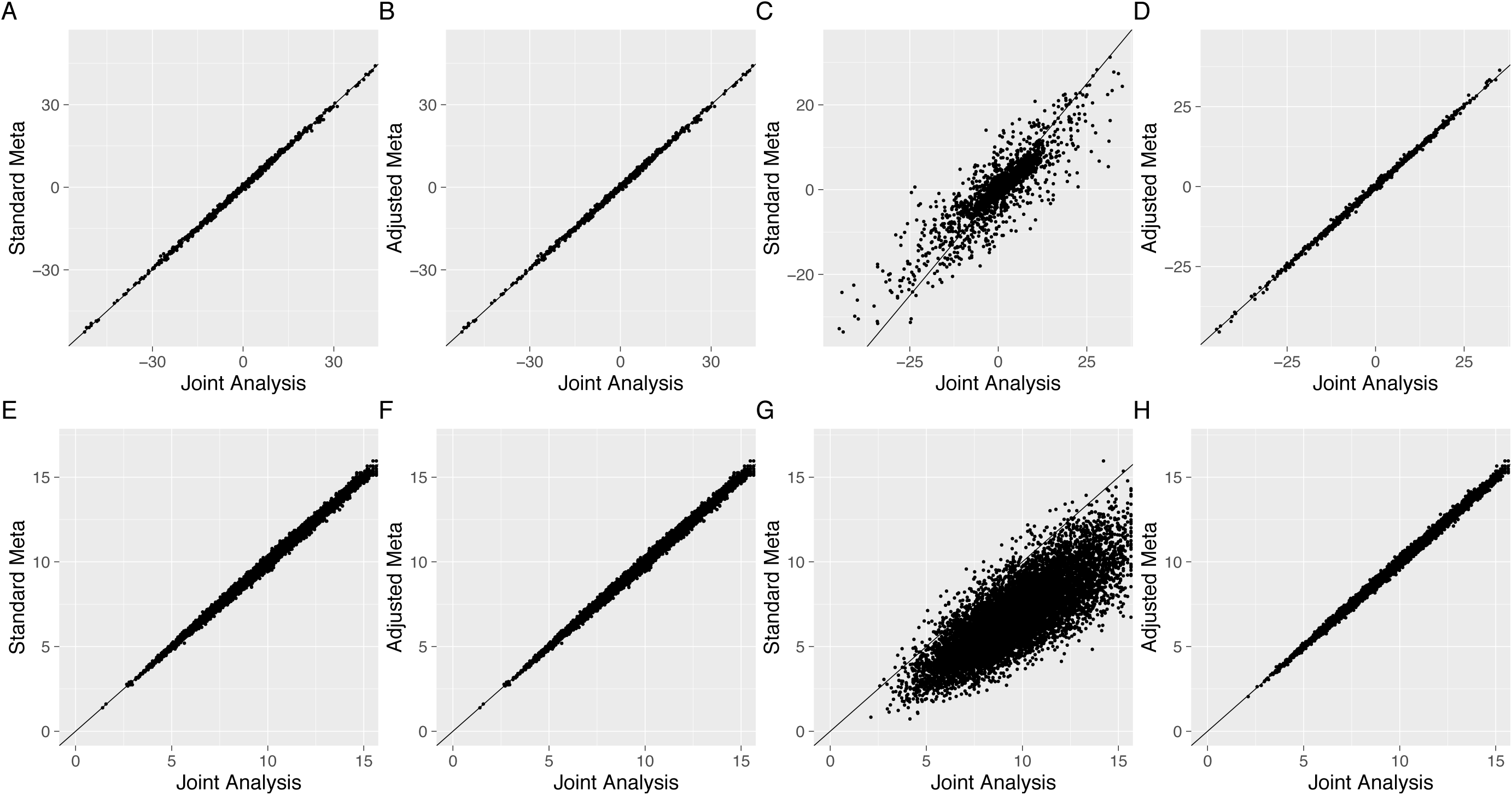
Score statistics (A, B, C, D) and – log10(p values) of the corresponding single-variant score tests (E, F, G, H), for dichotomous studies with common covariates, without population stratification, under balanced and unbalanced settings. (A, B): Score statistics under balanced studies; (C, D) Score statistics under unbalanced studies; (E, F): – log10 (p values) of single-variant score tests under balanced studies; (G, H) – log10 (p values) of single-variant score tests under unbalanced studies. “Standard Meta” denotes the standard meta-analysis methods; “Adjusted Meta” denotes our new meta-analysis methods.

With more accurate estimation for the joint score statistics, the gene-level tests (i.e., Burden test and SKAT based on score statistics) by our adjusted meta-analysis method are equivalent to joint analyses. This is the fundamental reason why our method performs as efficiently as the joint analysis under general settings, recovering up to 69% power loss caused by the standard method in unbalanced dichotomous studies with common covariates (Figure 3). Similar results were obtained for scenarios with different covariates (Figures S6-S8). Take the dichotomous studies with common covariates for examples (Figure 3), the power by standard meta-analysis method was 0.701 for Burden and 0.219 for SKAT, which were 27% and 69% less than the golden standards (0.964 for Burden; 0.703 for SKAT) by joint analysis (without population stratification); while the results by our adjusted meta-analysis method (power 0.964 for Burden; 0.702 for SKAT) were concordant with the joint analysis results.

**Figure 3.**
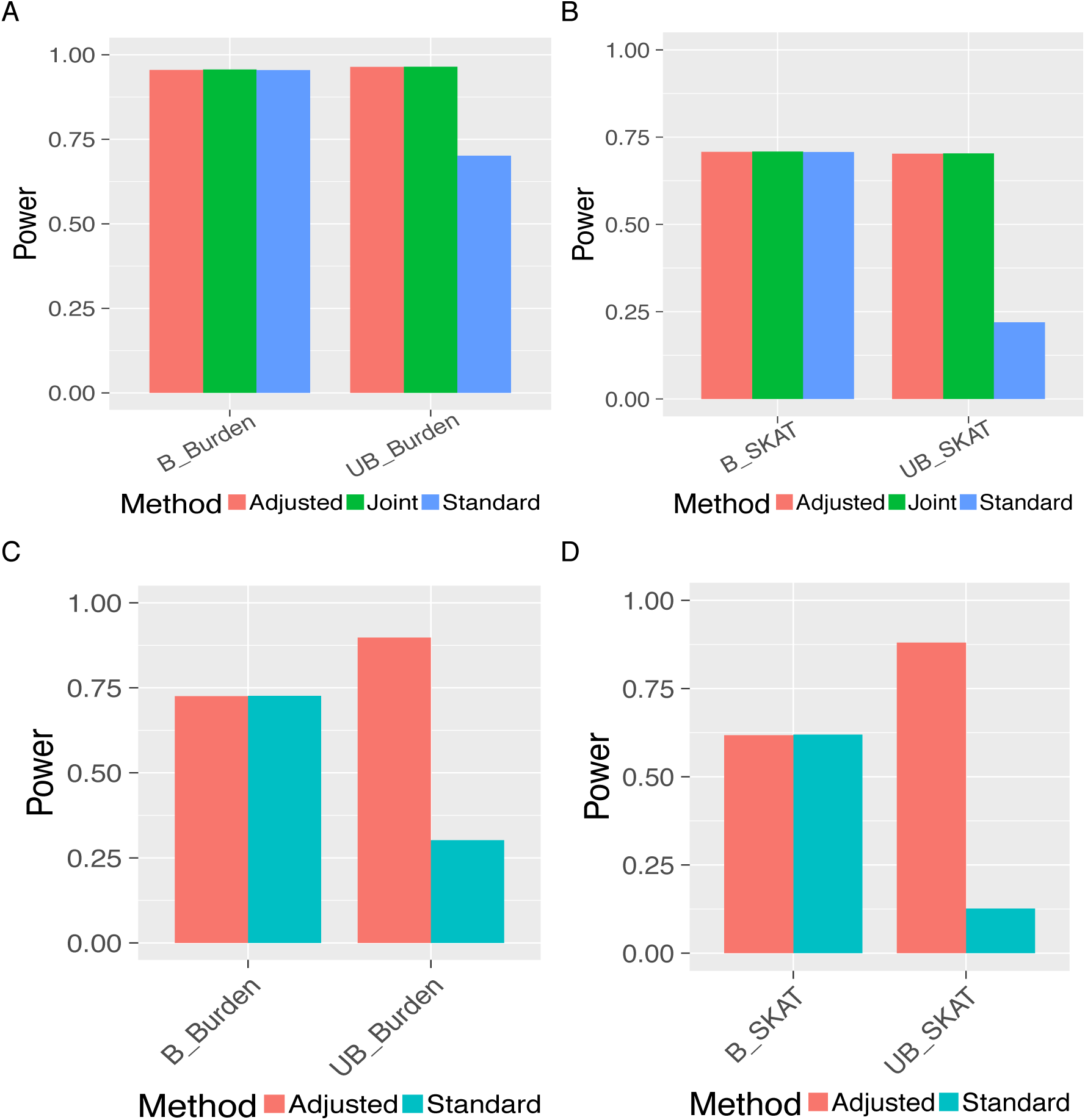
Power comparisons of meta-Burden test and meta-SKAT, for balanced (B) and unbalanced (UB) dichotomous studies with common covariates. (A, B): Without population stratification; (C, D): With population stratification. “Adjusted” denotes our new meta-analysis methods; “Standard” denotes the standard meta-analysis methods; and “Joint” denotes the joint analyses using combined individual-level data.

In the scenarios with population stratification, the joint analysis (with top 4 PCs as additional covariates) no longer provide golden standards due to highly inflated type I errors (see QQ plots of – log10 (p values) in Figure S5(A, B)). Hence, we only compared the empirical powers by our adjusted meta-analysis method with the standard method. Again, both methods had similar power in balanced dichotomous studies, while our adjusted meta-analysis method recovered up to 85% power loss by the standard method in unbalanced dichotomous studies (0.898 vs. 0.302 for Burden test, Figure 3C; 0.880 vs. 0.126 for SKAT, Figure 3D).

For quantitative studies, although we simulated “unbalanced” scenarios with various sample sizes, these are not really unbalanced for having about the same phenotype means across individual studies (i.e., the between study variances were close to 0). As a result, both our adjusted method and the standard method produced equivalent results as joint analyses under all settings (Figure S9-S13).

In summary, the simulations showed that our adjusted meta-analysis method will improve power by correctly modeling the association information in the between-study variances. When the between-study variances are close to 0 as under balanced settings, both our method and the standard method are equivalent to the joint analysis. When the between-study variances are also subject to population stratification, our method require good reference panels to correct for possibly inflated type I errors.

### Real Study of AMD

We applied our method on the real AMD data collected by the International AMD Genomics Consortium (IAMDGC)^23^, which has 26 individual studies with 33,976 European, 1,572 Asian, and 413 African unrelated samples. We treated the Asian and African samples as two extra studies. First, we conducted null simulations for 2.5 × 10^7^ times using the AMD data, by permuting the real AMD phenotypes and randomly selecting genotype regions of 100 variants for Burden test and SKAT. We found that both our adjusted and the standard meta-analysis methods controlled well for type I errors, while joint analyses with first 4 joint PCs as extra covariates resulted inflated type I errors (Figure S14). Specifically, with significance level 2.5 × 10^−6^, the joint analyses (Joint_PC4) had type I errors 8.6 × 10^−6^ for Burden test and 9.2 × 10^−6^ for SKAT.

For valid comparisons with joint analyses, we only considered European samples from the 26 unbalanced studies (Table S1) for Burden test and SKAT in 3 example AMD risk genes^23^ (*CFH*, *CFI*, *TIMP3*). Previous analyses by variable-threshold tests^27^ (with respective MAF thresholds 0.015%, 0.068%, 0.021% for genes *CFH*, *CFI*, *TIMP3*) gave significant p values (< 2.5 × 10^−6^) for these 3 loci. To be consistent with the previous variable-threshold tests^27^, we only analyzed protein-altering variants (imputed / genotyped) with MAFs under the corresponding thresholds (MAFs < 0.015%, 0.068%, 0.021%), and corrected for the same covariates ––– known independent signals within the same locus, gender, first two principal components (calculated using the combined data), and source of DNA (whole-blood or whole genome-amplified DNA).

Our adjusted meta-analysis method produced genome-wide significant p values for genes *CFH* and *CFI* (Table 1), which were more significant than the ones by the standard method. Specifically, gene *CFH* had genome-wide significant Burden p value 2.4 × 10^−7^ by joint analysis, versus 2.1 × 10^−6^ by our adjusted meta-analysis method and 3.2 × 10^−5^ by the standard method (no longer genome-wide significant). Although all methods obtained significant Burden p values for gene *CFI*, the p value by our method was still more significant than the one by the standard method (3.3 × 10^−14^ vs. 9.6 × 10^−10^) and closer to the p value by joint analysis (8.9 × 10^−15^). Similarly, the SKAT p value by standard method for gene *CFI* was no longer genome-wide significant (1.2 × 10^−4^), while the SKAT p value by our adjusted method (3.1 × 10^−9^) was still genome-wide significant and close to the one by joint analysis (1.9 × 10^−10^).

**Table 1.** P values of gene-level Burden test and SKAT by the standard meta-analysis method, our adjusted meta-analysis method, and joint analysis.

| Gene | Burden Test |  |  | SKAT |  |  |
| --- | --- | --- | --- | --- | --- | --- |
|  | Standard | Adjusted | Joint | Standard | Adjusted | Joint |
| CFH | $3.2 \times 10^{-5}$ | $2.1 \times 10^{-6}$ | $2.4 \times 10^{-7}$ | $6.1 \times 10^{-4}$ | $8.4 \times 10^{-5}$ | $3.6 \times 10^{-5}$ |
| CFI | $9.6 \times 10^{-10}$ | $3.3 \times 10^{-14}$ | $8.9 \times 10^{-15}$ | $1.2 \times 10^{-4}$ | $3.1 \times 10^{-9}$ | $1.9 \times 10^{-10}$ |
| TIMP3 | $9.8 \times 10^{-4}$ | $1.0 \times 10^{-5}$ | $1.8 \times 10^{-5}$ | $2.6 \times 10^{-3}$ | $7.4 \times 10^{-5}$ | $2.6 \times 10^{-4}$ |

Even though all approaches failed to identify the *TIMP3* locus with p values 1.8 × 10^−5^ by joint Burden test and 2.6 × 10^−4^ by joint SKAT, our method still produced more significant p values than the standard method (1.0 × 10^−5^ vs. 9.8 × 10^−4^ for Burden test; 7.4 × 10^−5^ vs. 2.6 ×10^−3^ for SKAT). Likely due to the meta-analysis errors, the Burden and SKAT p values for gene *TIMP3* by our method are slightly smaller than the ones by joint analysis.

This real example of AMD study demonstrated that both our method and the standard method controlled well for type I errors, and that our method outperformed the standard method by correctly modeling the between-study variances under unbalanced settings. Here, our method achieved the same power as the joint analysis whose results serve as golden standards with single-ethnic samples.

### Real Study of T2D

In this real example, we considered single-variant meta-analyses of three T2D GWASs: FUSION (1,142 cases vs. 1,155 controls; unrelated Finnish samples)^2^, METSIM (673 cases vs. 2,667 controls; unrelated Finnish male samples)^26^, and MGI (1,942 cases vs. 14,553 controls; unrelated European American samples). These three unbalanced GWASs have various case-control ratios (0.98, 0.24, 0.13) and multi-ethnic samples (Figures S15 and S16).

We first jointly corrected the T2D phenotypes for age, gender, body mass index (BMI), and first two joint PCs, within individual studies. The reason of jointly correcting the T2D phenotypes is to eliminate the possible between-study variance due to the artificial effects caused by individually corrected phenotypes. Then we applied the joint analysis, the standard, our joint-equivalent method (without adjustment for population stratification), and our adjusted meta-analysis method with adjustment for population stratification using the population-specific MAFs of EUR, AMR, AFR, SAS, EAS from the 1000 Genome Project^17^ (∽ 500 samples per population). In the step of adjusting for population by regressing known population-specific effects (MAFs) out from the within-study MAFs, the regression R^2^ was 97.1%, 96.3%, and 99.5% for FUSION, METSIM, and MGI studies, respectively. This showed that the 1000 Genome^17^ might not be the best reference panel for the FUSION and METSIM studies with Finnish samples, as > 99% regression R^2^ is expected for a good reference panel.

In this study, we only analyzed 631,870 variants that were genotyped in the METSIM study as an example. These analyzed variants could be either genotyped or imputed to 1000 Genome Project^17^ or absent in FUSION (627,920 variants) and MGI (631,628 variants) studies (see Manhattan plots of the individual GWASs in Figure S17). As expected, the joint analysis and the joint-equivalent methods resulted inflated type I errors (with inflated genomic control factors, λ_GC_ = 1.11, 1.13), because the between-study variances were also subject to population stratification (see QQ plots in Figure S18 (A, C)). The standard meta-analysis method was not affected by the population stratification for not modeling the between-study variances (λ_GC_ = 1.07, Figure S18 (B)). Specifically, the standard method identified three known T2D risk loci (*CDKAL1* on CHR6, *SLC30A8* on CHR8, and *TCF7L2* on CHR10)^28^, while our method with adjustment for population stratification identified comparable p values for signals in the *SLC30A8* and *TCF7L2* loci, more significant p value in the *CDKAL1* locus, and one extra potential loci *ROBO2* on CHR3 (see Manhattan plots in Figure 4).

**Figure 4.**
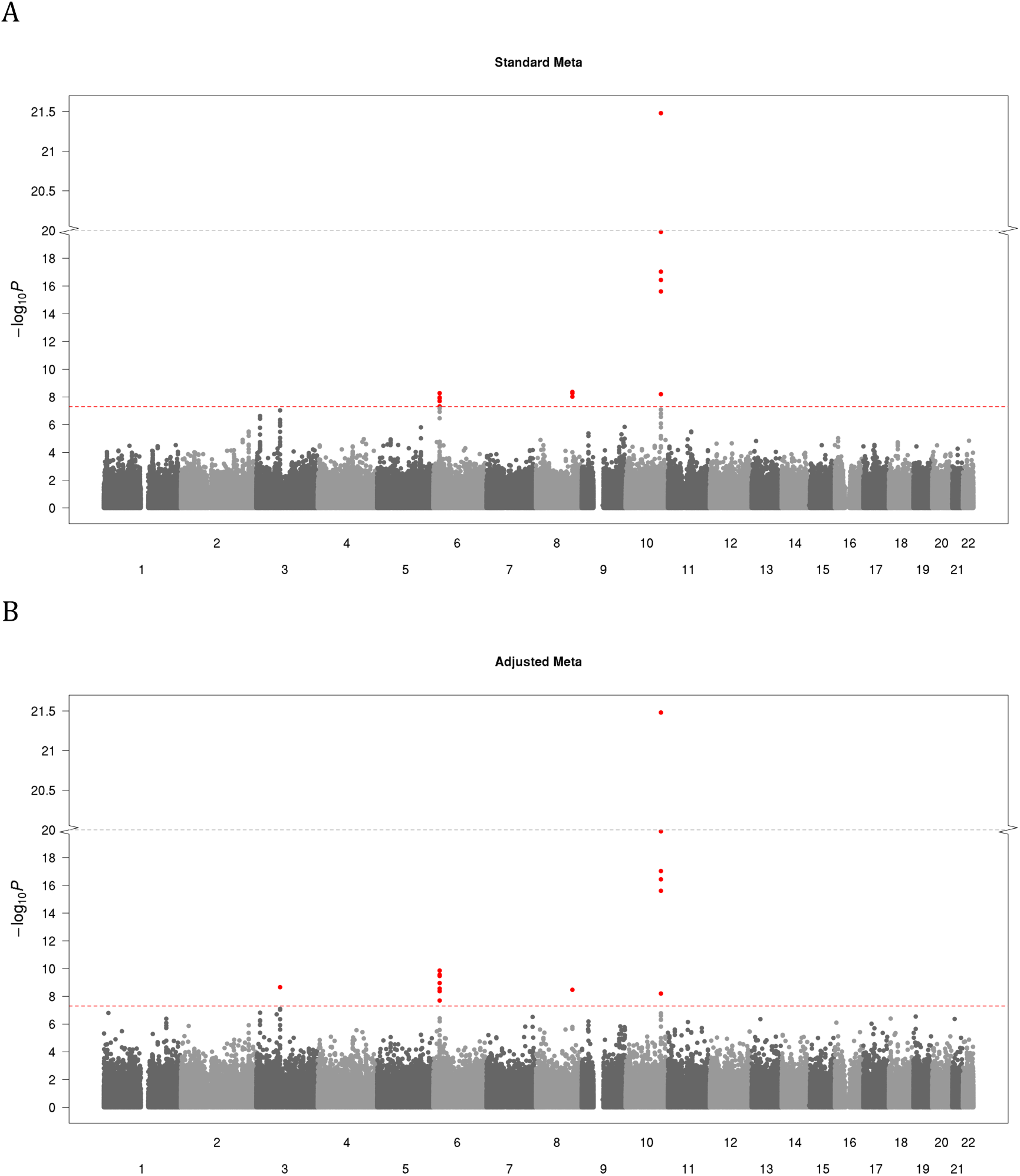
Manhattan plots of meta GWASs of type 2 diabetes, by standard method (A) and our adjusted method (B). “Standard Meta” denotes the standard meta-analysis methods; “Adjusted Meta” denotes our new meta-analysis methods.

We looked into the within-study MAFs of all “genome-wide significant” variants that were identified by joint analysis (Figure S19). We found that all “false positive” signals by joint analysis were likely due to the big differences among the within-study MAFs, while the “true” signals identified by the standard method have comparable within-study MAFs. For variants whose population-specific MAFs are “accurate” from the 1000 Genome Project^17^, our adjusted method could correct for the population stratification (i.e., MAF variations due to population differences). However, if the within-study MAFs were different due to mislabeled minor alleles, genotype errors, and small sample sizes, our adjusting step is likely to fail. The “inaccurate” population-specific MAFs from the reference panel and possible “wrong” within-study MAFs could potentially cause the “inflated” genomic control factor *λ_GC_* = 1.15 with our adjusted method (Figure S18 (D)).

This real study demonstrated the benefit of improving power by applying our adjusted meta-analysis method on unbalanced studies. Further, this study showed the challenges of correctly adjusting for population stratification when samples are of multi-ethnic. Our method requires “accurate” within-study MAFs and “accurate” population-specific MAFs from the reference panels. For cases where the adjustment of population stratification is likely to fail, we suggest using the standard method to be conservative.

## Discussion

In this paper, we propose improved formulas for accurately estimating the joint score statistics in meta-analysis, which had R^2^ > 99% with the ones obtainable using individual-level data under general settings. Consequently, for both single-variant score tests and gene-level tests based on score statistics (e.g., Burden test and SKAT), our meta-analysis method performs equivalently as the joint analysis using individual-level data under general settings. Importantly, our method is applicable for both linear and logistic regression models, with and without covariates. Both simulations and the real example of AMD demonstrated that our method performed as efficient as the joint analysis with unique-ethnic samples, substantially improving power over the standard method in unbalanced studies with various case-control ratios.

We further propose a novel approach to adjust for population stratification when the combined samples are of multi-ethnics. Observing that the population stratification is reflected by the differences of within-study MAFs in the score statistic formulas, we propose to normalize population structures by regressing out the effects of known population-specific MAFs (obtainable from external reference panels, e.g., 1000 Genome Project^17^ and Biobanks) from the within-study MAFs. Simulation studies with “accurate” population-specific MAFs based on 10^5^ samples showed the success of adjusting for population stratification by our adjusted meta-analysis method. This approach even avoids the dilemma of choosing an appropriate number of PCs as additional covariates. Both simulation and real studies demonstrated that our adjusted meta-analysis method controlled well for type I errors in general scenarios and gained power under unbalanced settings.

However, there are limitations about our method. First, our method assumes that the genetic effects are homogeneous across studies and the phenotypes are of the same distribution. Second, our method requires that there are no artificial effects involved in the between-study variances. Third, our method requires “accurate” within-study MAFs and good reference panel for correctly adjusting for population stratification. When the between-study variances contain information due to artificial effects or population stratification, the standard method is preferred for avoiding inflated false positive rates. Taking the real study of T2D as an example, we discussed the challenges of correctly adjusting for population stratification in practice.

In conclusion, we provide improved score statistic formulas in terms of summary statistics, for the analogous ones in joint analysis. These score statistics can then be used to conduct both single-variant and gene-level associations studies. Through these formulas, we showed that the between-study variances subject to population stratification (various MAFs across populations) are likely to cause inflated type I errors, and explained why the standard method is free of effects from population stratification. We further proposed a novel approach for adjusting population stratification using known population-specific MAFs from reference panels. As a result, our meta-analysis approach provides a useful framework ensuring well-powered, convenient, cross-study association analyses.

## Acknowledgments

This work was supported by the National Institutes of Health (NIH) grant R01HG007022. All authors claim there is no conflict of interest. The authors would like to thank Dr. Dajiang Liu (Pennsylvania State University), Dr. Shuang Feng (graduated Ph.D. student from University of Michigan), and Dr. Shawn Lee (University of Michigan) for their inspiration and valuable comments for this work. Especially, the authors want to thank all investigators of the IAMDGC, FUSION, METSIM, and MGI studies for providing the real data of the AMD and T2D studies in this work.

## Web Resources

Software RAREMETAL, https://github.com/traxexx/Raremetal.

## Appendices

### Appendix A: Score Statistics for an individual Study

For the *k* th sub-study, denote the mean genotype matrix by 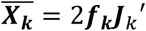, where *f*_*k*_ denotes the within-study MAF vector and *J*_*k*_ denotes a *n*_*k*_ × 1 vector of 1’s. The score statistic vector and corresponding variance-covariance matrix are given by

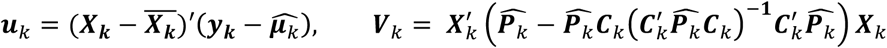

where 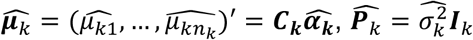 for quantitative traits under the standard linear regression model; 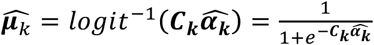, 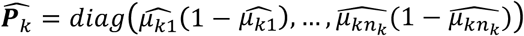 for dichotomous traits under the standard logistic regression model; coefficient vector 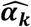 and residual variance 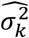 are estimated under the null model *β*_*k*_ = 0; and ***I***_*k*_ denotes a *n_k_ × n_k_* identity matrix.

### Appendix B: Score Statistics for Combined Data

For simplicity of notations, assume all *K* studies have the same set of genetic variants and covariates. Let 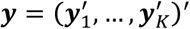 denotes the joint n × 1 phenotype vector of *K* studies, 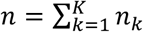, 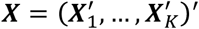 denotes the joint n × m genotype matrix, and 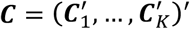 denotes the joint n × (q + 1) augmented covariate matrix. Denote the overall mean genotype matrix by 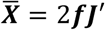, where ***f*** is the overall MAF vector and ***j*** is a n × 1 vector of 1’s. With combined data (**X, y, C**), the joint score statistic vector and corresponding variance-covariance matrix are given by

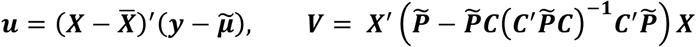

where 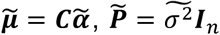 for quantitative traits under the standard linear regression model; 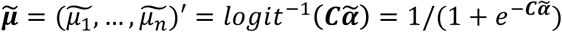, 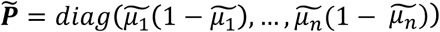 for dichotomous traits under the standard logistic regression model; coefficient vector 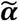 and residual variance 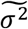 are estimated under the null model (β= 0); ***I***_n_ denotes a n × n identity matrix.

### Appendix B.1: Score Statistics for the Simplified Case without Covariates

Without covariates, for both quantitative and dichotomous traits, we have within-study phenotype mean 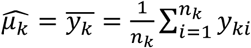; overall phenotype mean 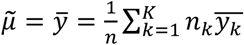; phenotype deviation 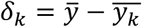; and 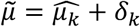. Note that 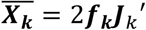 and 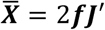. The joint score statistic vector is given by

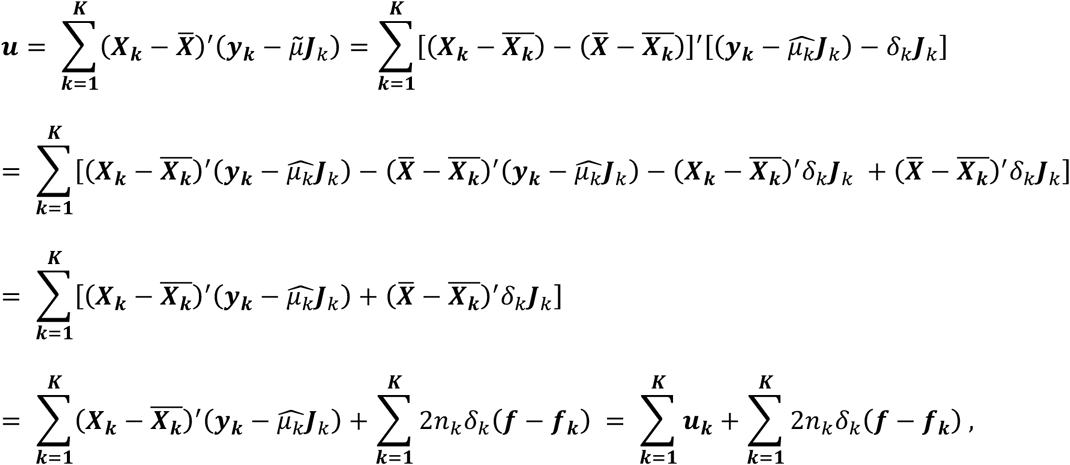

where 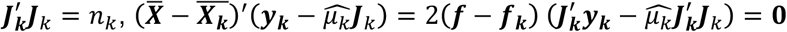 and 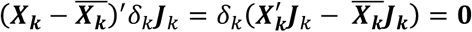

Because *C*_k_ = *J*_***k***_ in the case without covariates, the covariance matrix of the score statistic can be simplified as

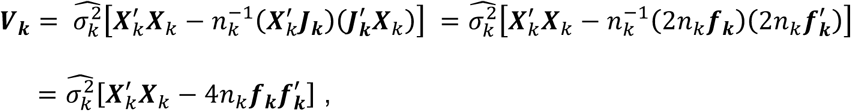

where 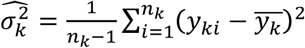 for quantitative traits under the standard linear regression model, and 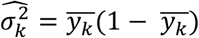 for dichotomous traits under the standard logistic regression model.

Similarly, the joint ***V*** is given by

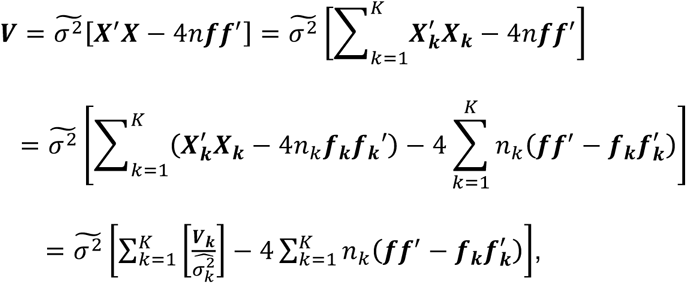

where 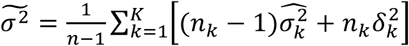 for quantitative traits under the standard linear regression model, and 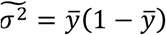 for dichotomous traits under the standard logistic regression model.

### Appendix B.2: Score Statistics with Covariates

In cases with covariates, the same formula is derived for approximating the joint score statistic vector ***u*** as in the simplified case (Appendix B.1) but with 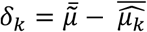, where 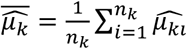 is the average of the fitted mean 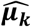 in the null model (study k); 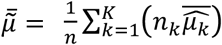 is the approximated average of the fitted mean 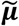 with combined data. Note that 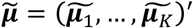, and 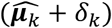 is an unbiased estimate for 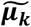. The formulas for the variance-covariance matrix ***V*** will be more complicated.

Under the standard linear regression model (Equation 1) for quantitative traits, 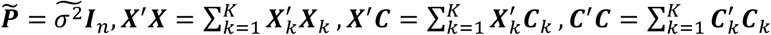. We can write ***V*** as

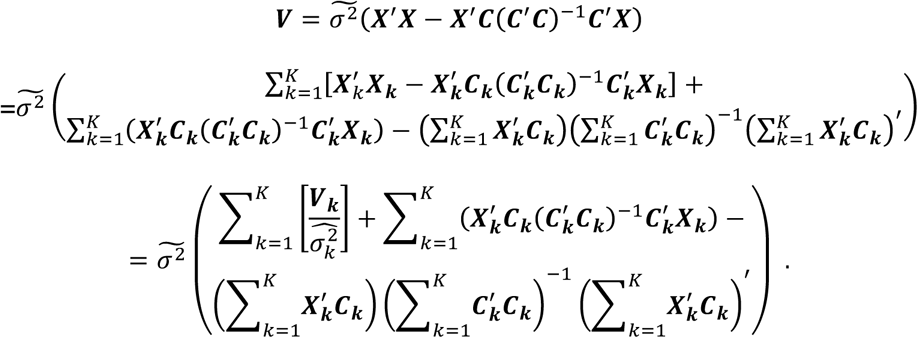

In this case, 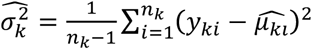 and 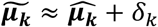the estimate of the noise variance in the joint model 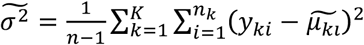 can be approximated by

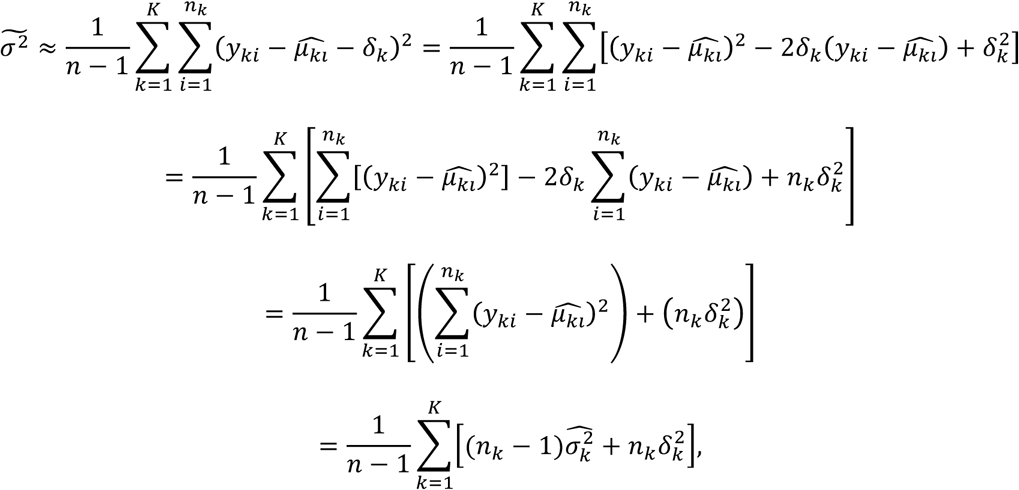

where 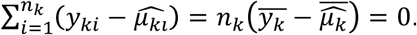.

Under the standard logistic regression model (Equation 2) for dichotomous traits, 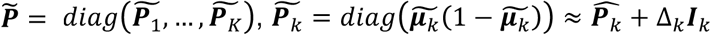 with

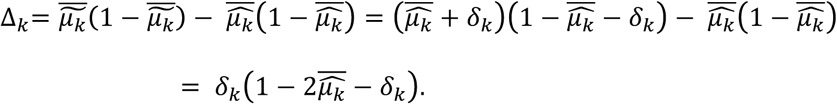

We can write *V* as

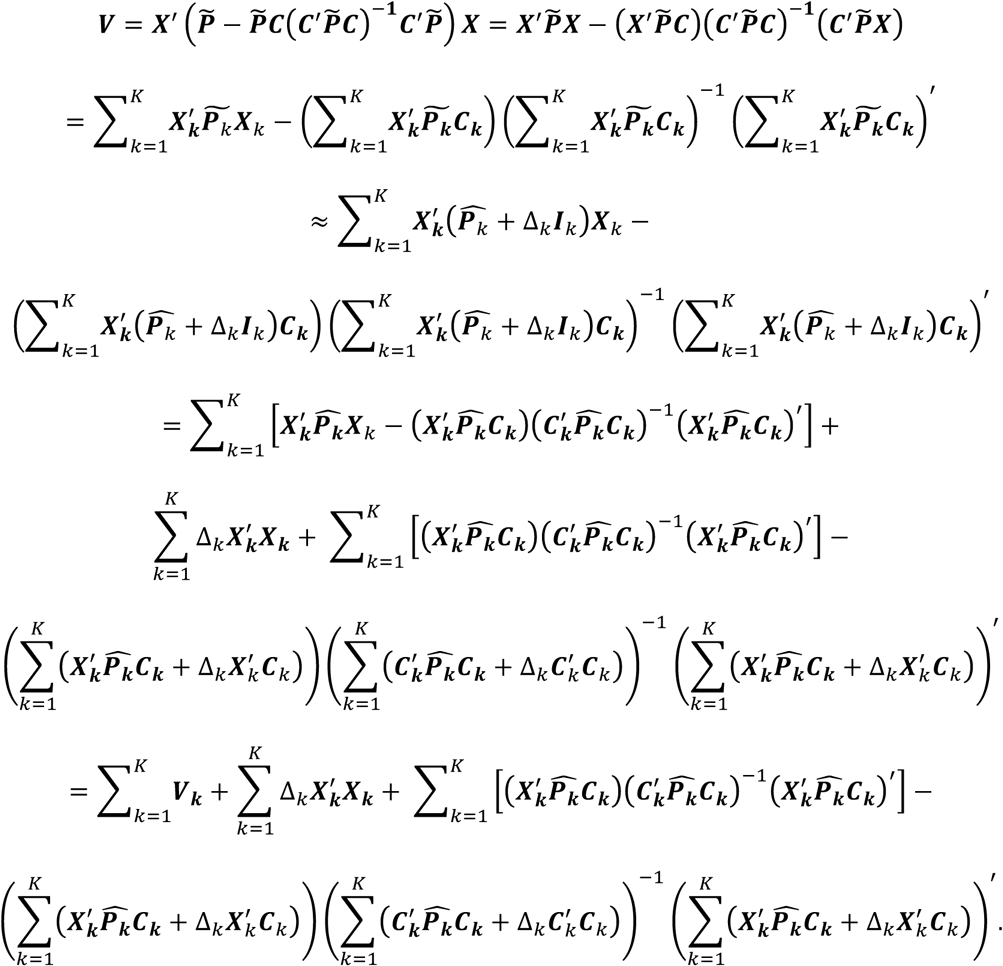

To use the above formulas to estimate ***V***, besides 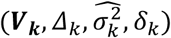, the quantities of 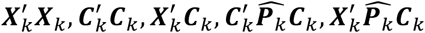 also need to be shared from individual studies.

### Appendix C: Simulation Studies

We first simulated a pool of 20,000 haplotypes (with length 5KB) for each of the three populations, European (EUR), Asian (ASA), and African (AFR), by COSI with the well calibrated coalescent model^24^. Then we sampled genotypes of 1×10^5^ individuals per population with 339 variants, where 96% are rare with MAFs < 5%. Regions of 100 variants were randomly selected as risk loci to simulate phenotypes (see spectrum plots of log10(MAFs) in Figure S1).

We simulated both quantitative and dichotomous phenotypes according to the respective standard linear and logistic models (Equations 1 and 2). Under the linear regression model (Equation 1), we selected residual variance 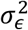 and effect-sizes β such that a given amount of heritability was equally explained by all causal variants; under the logistic model (Equation 2), we selected the intercept term subject to 1% disease prevalence and the log-odds-ratios *β* by 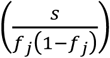 with a given constant *s*. These parameters will be set to result in ∽ 80% power in the simulations, and equal variance contributions from all causal variants.

Two covariate scenarios were simulated: (i) common covariates (*C*_1_, *C*_2_) for all studies; (ii) different covariates among studies –– the first and third studies have single covariate (*C*_1_), the third study has two covariates (*C*_1_, *C*_2_), while the second and fifth studies have three covariates (*C*_1_, *C*_2_, *C*_3_). Specifically, *C*_1_ is a binary covariate generated from *Bernoulli (p = 0.5)*; *C*_2_ and *C*_3_ are continuous covariates generated from *N*(0, 1). The covariate coefficients were selected such that 1% phenotype variance was equally explained per covariate in the linear model, i.e., (*α*_1_ = 0.2), (*α*_1_ = 0.141, *α*_2_ = 0.071), (*α*_1_ = 0.115, *α*_2_ *α*= _3_ = 0.057), respectively for models with covariates (*C*_1_), (*C*_1_, *C*_2_), *C*_1_, *C*_2_, *C*_3_. The covariates coefficients were taken as 0.1 in the logistic model.

For each scenario, we first generated 100 sets of phenotypes and corresponding covariates for all samples in the population. Then we randomly drew samples with corresponding phenotypes, genotypes, and covariates from the simulated populations.

## References

1. Lin, D.Y., and Zeng, D. (2010). Meta-analysis of genome-wide association studies: no efficiency gain in using individual participant data. Genetic epidemiology 34, 60 – 66.

2. Scott, L.J., Mohlke, K.L., Bonnycastle, L.L., Willer, C.J., Li, Y., Duren, W.L., Erdos, M.R., Stringham, H.M., Chines, P.S., Jackson, A.U., et al. (2007). A genome-wide association study of type 2 diabetes in Finns detects multiple susceptibility variants. Science 316, 1341 – 1345.

3. Zeggini, E., Scott, L.J., Saxena, R., Voight, B.F., Marchini, J.L., Hu, T., de Bakker, P.I., Abecasis, G.R., Almgren, P., Andersen, G., et al. (2008). Meta-analysis of genome-wide association data and large-scale replication identifies additional susceptibility loci for type 2 diabetes. Nature genetics 40, 638 – 645.

4. Fuchsberger, C., Flannick, J., Teslovich, T.M., Mahajan, A., Agarwala, V., Gaulton, K.J., Ma, C., Fontanillas, P., Moutsianas, L., McCarthy, D.J., et al. (2016). The genetic architecture of type 2 diabetes. Nature 536, 41 – 47.

5. Willer, C.J., Sanna, S., Jackson, A.U., Scuteri, A., Bonnycastle, L.L., Clarke, R., Heath, S.C., Timpson, N.J., Najjar, S.S., Stringham, H.M., et al. (2008). Newly identified loci that influence lipid concentrations and risk of coronary artery disease. Nature genetics 40, 161 – 169.

6. Willer, C.J., Speliotes, E.K., Loos, R.J., Li, S., Lindgren, C.M., Heid, I.M., Berndt, S.I., Elliott, A.L., Jackson, A.U., Lamina, C., et al. (2009). Six new loci associated with body mass index highlight a neuronal influence on body weight regulation. Nature genetics 41, 25 – 34.

7. Stahl, E.A., Raychaudhuri, S., Remmers, E.F., Xie, G., Eyre, S., Thomson, B.P., Li, Y., Kurreeman, F.A., Zhernakova, A., Hinks, A., et al. (2010). Genome-wide association study meta-analysis identifies seven new rheumatoid arthritis risk loci. Nature genetics 42, 508 – 514.

8. Prokopenko, I., Langenberg, C., Florez, J.C., Saxena, R., Soranzo, N., Thorleifsson, G., Loos, R.J., Manning, A.K., Jackson, A.U., Aulchenko, Y., et al. (2009). Variants in MTNR1B influence fasting glucose levels. Nature genetics 41, 77 – 81.

9. Willer, C.J., Li, Y., and Abecasis, G.R. (2010). METAL: fast and efficient meta-analysis of genomewide association scans. Bioinformatics 26, 2190 – 2191.

10. Lee, S., Teslovich, T.M., Boehnke, M., and Lin, X. (2013). General framework for meta-analysis of rare variants in sequencing association studies. American journal of human genetics 93, 42 – 53.

11. Tang, Z.Z., and Lin, D.Y. (2013). MASS: meta-analysis of score statistics for sequencing studies. Bioinformatics 29, 1803 – 1805.

12. Feng, S., Liu, D., Zhan, X., Wing, M.K., and Abecasis, G.R. (2014). RAREMETAL: fast and powerful meta-analysis for rare variants. Bioinformatics 30, 2828 – 2829.

13. Liu, D.J., Peloso, G.M., Zhan, X., Holmen, O.L., Zawistowski, M., Feng, S., Nikpay, M., Auer, P.L., Goel, A., Zhang, H., et al. (2014). Meta-analysis of gene-level tests for rare variant association. Nature genetics 46, 200 – 204.

14. Sudlow, C., Gallacher, J., Allen, N., Beral, V., Burton, P., Danesh, J., Downey, P., Elliott, P., Green, J., Landray, M., et al. (2015). UK biobank: an open access resource for identifying the causes of a wide range of complex diseases of middle and old age. PLoS Med 12, e1001779.

15. Stouffer, S.A.S., Edward A; DeVinney, Leland C; Star, Shirley A; Williams Jr, Robin M. (1949). The American soldier: adjustment during army life. (Oxford, England: Princeton University Press).

16. Cochran, W.G. (1954). The combination of estimates from different experiments. Biometrics 10, 101–129.

17. Genomes Project, C., Abecasis, G.R., Auton, A., Brooks, L.D., DePristo, M.A., Durbin, R.M., Handsaker, R.E., Kang, H.M., Marth, G.T., and McVean, G.A. (2012). An integrated map of genetic variation from 1,092 human genomes. Nature 491, 56–65.|

18. Price, A.L., Patterson, N.J., Plenge, R.M., Weinblatt, M.E., Shadick, N.A., and Reich, D. (2006). Principal components analysis corrects for stratification in genome-wide association studies. Nature genetics 38, 904–909.

19. Radhakrishna Rao, C. (1948). Large sample tests of statistical hypotheses concerning several parameters with applications to problems of estimation. Mathematical Proceedings of the Cambridge Philosophical Society 44, 50–57.

20. Morris, A.P., and Zeggini, E. (2010). An evaluation of statistical approaches to rare variant analysis in genetic association studies. Genetic epidemiology 34, 188–193.

21. Neale, B.M., Rivas, M.A., Voight, B.F., Altshuler, D., Devlin, B., Orho-Melander, M., Kathiresan, S., Purcell, S.M., Roeder, K., and Daly, M.J. (2011). Testing for an unusual distribution of rare variants. PLoS genetics 7, e1001322.

22. Wu, M.C., Lee, S., Cai, T., Li, Y., Boehnke, M., and Lin, X. (2011). Rare-variant association testing for sequencing data with the sequence kernel association test. American journal of human genetics 89, 82 – 93.

23. Fritsche, L.G., Igl, W., Bailey, J.N., Grassmann, F., Sengupta, S., Bragg-Gresham, J.L., Burdon, K.P., Hebbring, S.J., Wen, C., Gorski, M., et al. (2015). A large genome-wide association study of age-related macular degeneration highlights contributions of rare and common variants. Nature genetics.

24. Schaffner, S.F., Foo, C., Gabriel, S., Reich, D., Daly, M.J., and Altshuler, D. (2005). Calibrating a coalescent simulation of human genome sequence variation. Genome research 15, 1576 – 1583.

25. Yang, J., Fritsche, L.G., Zhou, X., Abecasis, G., and International Age-Related Macular Degeneration Genomics, C. (2017). A Scalable Bayesian Method for Integrating Functional Information in Genome-wide Association Studies. American journal of human genetics.

26. Laakso, M., Kuusisto, J., Stancakova, A., Kuulasmaa, T., Pajukanta, P., Lusis, A.J., Collins, F.S., Mohlke, K.L., and Boehnke, M. (2017). The Metabolic Syndrome in Men study: a resource for studies of metabolic and cardiovascular diseases. Journal of lipid research 58, 481 – 493.

27. Price, A.L., Kryukov, G.V., de Bakker, P.I., Purcell, S.M., Staples, J., Wei, L.J., and Sunyaev, S.R. (2010). Pooled association tests for rare variants in exon-resequencing studies. American journal of human genetics 86, 832 – 838.

28. Billings, L.K., and Florez, J.C. (2010). The genetics of type 2 diabetes: what have we learned from GWAS? Ann N Y Acad Sci 1212, 59 – 77.

